# The turbulent brain: Modelling vortex interactions for understanding human cognition

**DOI:** 10.1101/2025.09.25.678538

**Authors:** Gustavo Deco, Yonatan Sanz Perl, Jianfeng Feng, Morten L. Kringelbach

## Abstract

The human brain needs distributed, time-critical computation to efficiently solve complex problems. Turbulence provides such highly efficient spacetime information processing and transmission across wide-spread brain networks, yet we have been missing a mechanistic understanding of the interactions of turbulent vortices underlying human cognition. Here, we build the first whole-brain model of turbulent vortices as defined by the levels of local synchronisation in brain signals quantifying turbulent interactions in vortex space. Specifically, using large-scale human neuroimaging data, we found that the interactions of turbulent vortices is an excellent framework for understanding cognition and brain computation. It is not only useful for distinguishing the detailed spacetime dynamics of rest and cognition – but can even distinguish between subtle sub-components of cognitive tasks, where manipulation of vortices can be shown to change cognition. Overall, this whole-brain framework creates a natural vortex space for the brain computation underlying cognition, as well as potentially providing novel ways of controlling turbulent interactions in disease.

## Introduction

In people with severe epilepsy, traumatic brain injury and post-traumatic stress disorder (PTSD), seemingly innocuous stimuli like certain smells can lead to a full-blown attack with potential catastrophic impact on the whole-brain dynamics (Hinojosa *et al*., 2024; Jirsa *et al*., 2023; Martinez-Molina *et al*., 2024). Equally, negative experiences can lead to rumination which can spiral out of control and become clinical depression (Figueroa *et al*., 2019). These forms of catastrophic impact are consistent with the role of turbulence as a potent form of spatiotemporal chaos that underlies phenomena like of how small perturbations, like the flapping of butterfly wings can potentially cause huge system-wide disturbance (Lorenz, 1993). In fact, recent research has demonstrated that the human brain is turbulent (Deco and Kringelbach, 2020; Deco *et al*., 2023; Deco *et al*., 2025). Furthermore, it has been demonstrated that measures of whole-brain turbulent dynamics are excellent for predicting the responsiveness to pharmacological treatment in major depressive disorder (Escrichs *et al*., 2024). Yet, we are still lacking an understanding of why this happens and how to ultimate control this. As such, a whole-brain modelling framework of turbulence found in brain dynamics could be a promising avenue for understanding the underlying dynamics and a first step on the road to controlling turbulence in disease.

Importantly, brain turbulence is not found in the fluids of the brain but rather in the local level of synchronisation of brain activity across space and time (Deco and Kringelbach, 2020; Deco *et al*., 2023; Kawamura *et al*., 2007; Kuramoto, 1984). This synchronisation generates turbulent vortices resembling the whirls from turbulent fluid dynamics (Frisch, 1995; Kolmogorov, 1941a, b) (Bewley *et al*., 2006; Christoph *et al*., 2018). Intuitively, any system that has high variability of this local level of synchronisation is turbulent and with this comes all the important properties needed for information transfer (Escrichs *et al*., 2022). Hence, a better understanding of the turbulent dynamics in the human brain could potentially lead to better ways of controlling the system from spinning out of control and become the basis of more effective treatments. Still, reaching this goal requires a deeper understanding of the interactions underlying turbulent dynamics in different brain states.

Here, we build the first whole-brain model of turbulent vortices to provide a mechanistic understanding of how their interactions underly the brain computation of human cognition. This can provide a precise description, which eventually could be controlled in much more efficient ways and perhaps become the basis of novel treatments.

Still, before such careful control can be implemented, a deeper understanding is needed of the interactions between turbulent vortices underlying normal states of cognition. As we show here, this provides a precise way to distinguish different cognitive tasks from rest, as well as distinguishing the more subtle sub-components of cognitive tasks. We also demonstrate that, in principle, manipulation at the level of turbulent vortices can change cognition, i.e., from resting to task. Overall, the findings demonstrate that the turbulent vortex space is a natural basis for the brain computation underlying cognition.

## Results

In order to model human cognition, many researchers have examined the fine-grained BOLD signals obtained with fMRI as they solve various tasks. However, using these BOLD timeseries is rarely sufficient for successfully decoding and distinguishing significant aspects of different tasks (Xu *et al*., 2023). Instead, based on the emergent literature on turbulence in brain dynamics, we came to the idea that a description of the interactions between turbulent vortices could provide a more direct way to model, understand and control brain dynamics. This intuition is based on the existing research on turbulence in fluid dynamics, where the vortex level provides a convenient basis for modelling, understanding and control.

**Figure 1** shows the overall schema of how the principles of turbulence have been firmly established in fluid dynamics at both microscopic and mesoscopic levels (**Figure 1A**), where power laws have been found that facilitate optimal mixing of the energy/information cascade. Importantly, while the microscopic signals can be described using the Navier-Stokes equations, these are not always computationally tractable. Instead, researchers use the interactions between macroscopic fluid vortices, which also allows them to understand the spacetime dynamics in a more transparent way and to control the levels of turbulence (Protas, 2008; Saffman, 1992).

**Figure 1.**
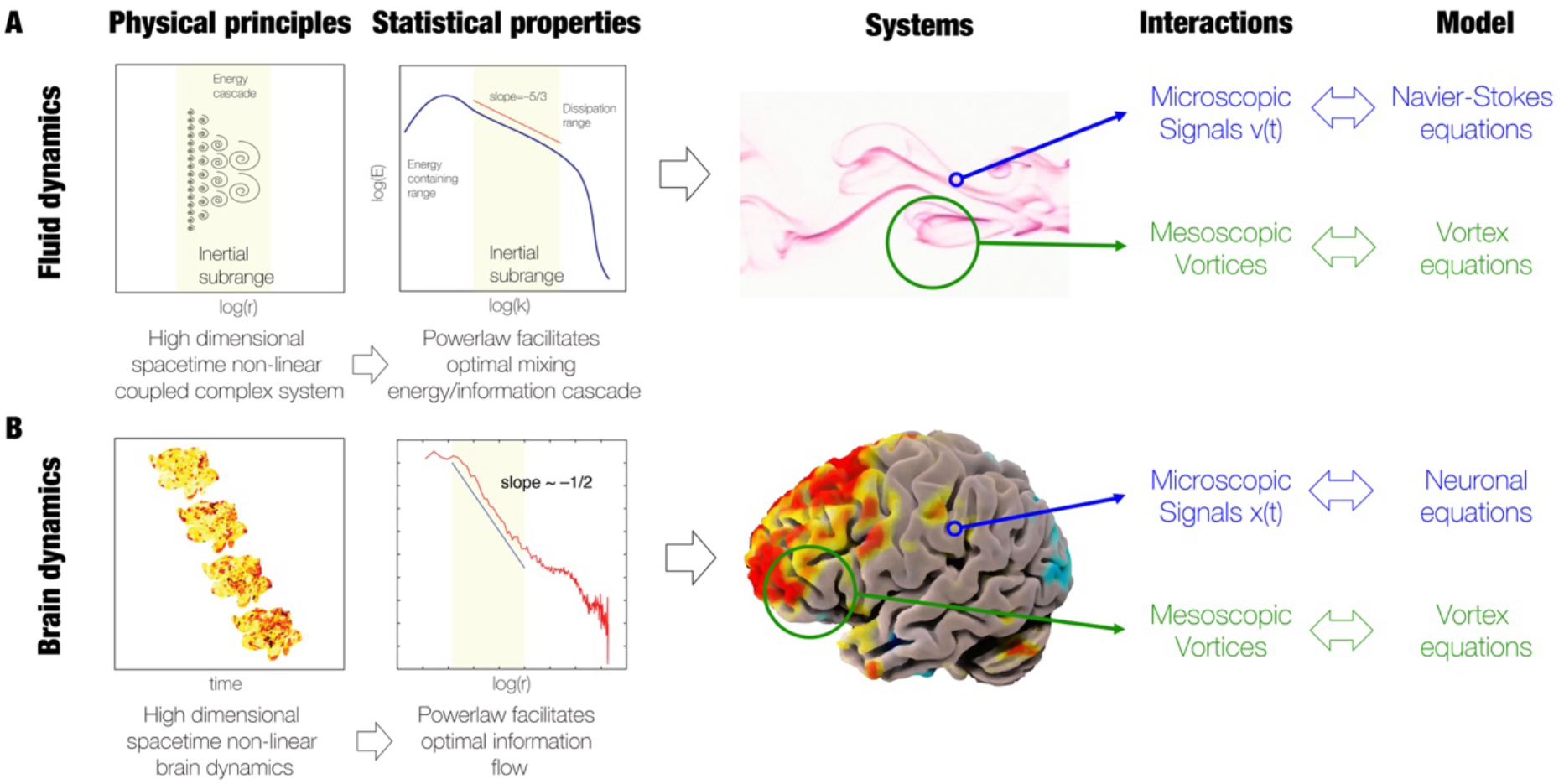
Overview of turbulence in fluids and brain oscillations. **A)** Following the discovery and naming of “turbolenza” by Leonardo da Vinci for the whirls of fluids, the science of turbulence was worked out over the next centuries. A significant advance was provided by Kolmogorov’s phenomenological theory of turbulence which is based on the concept of structure functions, inspired by Richardson’s concept of cascading eddies. This describes the statistical properties of the high dimensional space non-linear fluid dynamics (left panel). Kolmogorov discovered power laws in an inertial subrange where the structure functions show a universal energy scaling of 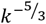, where k is the associated wave number of the spectral scale (middle panel). This power law behaviour reflects the energy/information transfer cascade found in turbulence. The right panel shows how turbulence can be modelled at the microscopic level by the Navier-Stokes equation but equally at the mesoscopic level by vortex equations, allowing a more efficient description and control of the dynamics. **B)** Turbulence is also found in non-fluid systems as demonstrated by Kuramoto, who used coupled-oscillators to describe the turbulent whirls of synchronised oscillators promoting optimal mixing. Turbulence is also found in brain dynamics (shown in the flattened brain renderings in left panel), where the phases of brain signal timeseries can be described over time and space by a local Kuramoto order parameter, reflecting the amplitude turbulence. This captures the evolution of the rich variability of the whirls of local synchronisation in brain dynamics. The middle panel shows how these turbulent dynamics also follow a power law, reflecting the optimality of efficiency of spacetime information flow in brain dynamics. Here we hypothesise that similar to the principles of fluid dynamics a mesoscopic description of the vortex space will provide a direct and more efficient way to describe and control brain dynamics (right panel).

Similarly, **Figure 1B** shows how turbulence can also be described using a system of coupled oscillators (Kuramoto, 1984). In particular, this form of turbulence has been found in brain dynamics (Deco and Kringelbach, 2020; Deco *et al*., 2023; Deco *et al*., 2025), where there are similar power laws (suggestive of fast information cascades) in the non-equilibrium brain dynamics (Perl *et al*., 2023a). As shown in the figure, the microscopic signals can be described by neuronal equations such as those established by Hodgkin-Huxley (Hodgkin and Huxley, 1952) but similar to the Navier-Stokes equations, these are also not always computationally tractable. Instead, here we propose to focus on the mesoscopic vortex space and use a whole-brain model to extract the interactions between turbulent vortices. Like the mesoscopic level in fluid dynamics, the mesoscopic level in oscillatory dynamics could offer a powerful basis for describing the distributed computation underlying cognition.

One potential avenue for describing the vortex space was provided by Xu and colleagues who extracted the phases and measured the curl of the flow of information in human neuroimaging data (Xu *et al*., 2023) (see **Figure 2**). They found that the resulting spiral-like, rotational wave patterns are widespread during both resting and cognitive task states, propagating across the cortex while rotating around their phase singularity centres, giving rise to spatiotemporal activity dynamics with non-stationary features. They demonstrated that these properties of brain spirals provide sensitive measures that can be used to classify different cognitive tasks.

**Figure 2.**
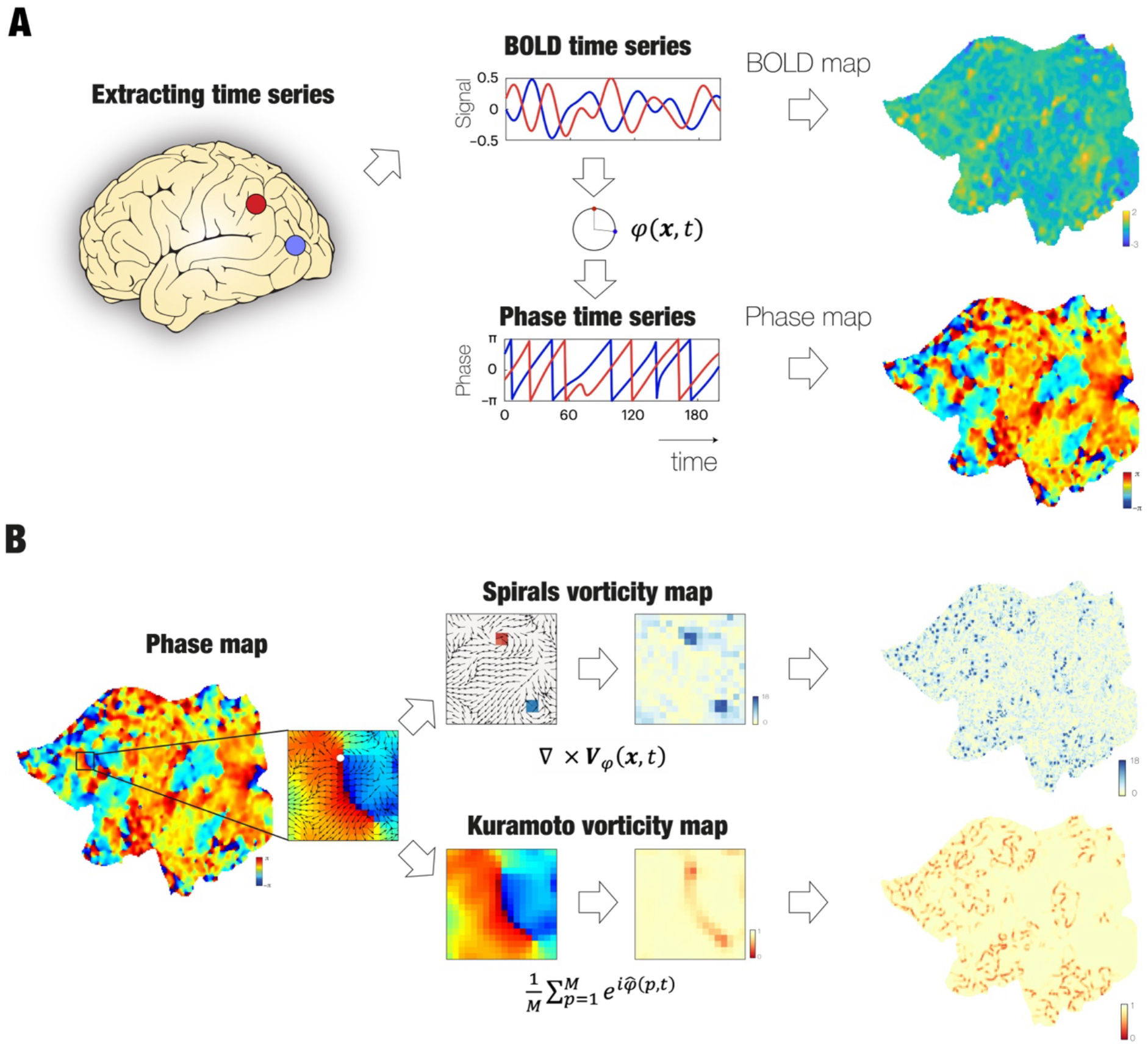
Pipeline for extracting vortices. **A)** Neuroimaging provides BOLD time series for each voxel in 3D space. Each voxel in 3D space is transformed to a flattened 2D cortical vertex space and the BOLD time series are transformed into phase time series by using the Hilbert transform. On the left is shown the extraction of the BOLD time series of two brain signals (blue and red, top panel) that are transformed into phase time series (bottom panel). At a given time point, this can be rendered on the flattened surface, where on the top right is shown snapshots of the BOLD values and bottom right the corresponding phase values. **B)** There are two ways of extracting vortices from the phase maps: The top middle panel shows the spiral vorticity map resulting from the curl of the flow of the phases. The bottom middle panel shows the Kuramoto vorticity map obtained by directly computing the local Kuramoto order parameters on the phases (see Methods). The surface renderings show a snapshot of this.

However, it is not easy to derive a model of brain spiral dynamics for quantifying their interactions. At the same time these brain spirals are intimately linked to turbulent vortices (Deco and Kringelbach, 2020; Deco *et al*., 2023; Deco *et al*., 2025) (see **Figure 2B**). Indeed, it is straightforward to build a model of coupled oscillators underlying turbulence, given that this research already uses the local Kuramoto order parameter (Kawamura *et al*., 2007) to measure the level of turbulence in oscillatory systems such as the brain (Deco and Kringelbach, 2020; Deco *et al*., 2023; Deco *et al*., 2025). In fact, the use of Kuramoto oscillators creates a natural interpretation for measuring the flow between turbulent vortices and, importantly, allowing for direct modelling of the interactions between vortices.

In a technical tour-de-force, here we created a vortex space from the fine-grained vertex space created by neuroimaging BOLD signals, where a vertex is the projection of a 3D brain voxel to the cortical surface. From this space, the interactions between each vertex can be reduced to a more coarse-grained parcellation of 1,000 regions (Schaefer1000) to an even more coarse-grained partition of 100 regions (Schaefer100). This is achieved using an influential partition technique (Snyder *et al*., 2020), effectively creating a lower dimensional vortex space, where an explicit equation can be derived for the modelling of the interactions between the 100 vortices.

This vortex space can be used as the basis of a whole-brain model which can quantify the interactions between vortices in vortex space. In other words, this framework is focusing on the vortex space rather than the signal vertex space obtained from BOLD measurements. This framework is thus moving away from modelling signal vertex space to modelling vortex space, which is crucial to capture the essential elements of the information cascade in turbulence. Excitingly (as shown in the *Methods*), modelling the interactions in vertex space with coupled Kuramoto local oscillators naturally leads to the modelling the vortex space with the celebrated Stuart-Landau equation (sometimes also called the Hopf equation), which is universally used in physics to measure most if not all physical phenomena. As such the research follows the dictum of Feynman who wrote “What I cannot create I do not understand” which emphasises the importance of modelling for understanding any physical phenomenon.

### Comparing Kuramoto vs spiral vorticity over time and space

As stated above, there is intuitively a close link between brain spirals and Kuramoto vorticity. In order to quantify this intuition, we directly compared the two (**Figure 3**). Following Xu and colleagues (Xu *et al*., 2023) – and as shown in the *Methods* – we used the definition of brain spirals in vertex space as the curl of the flow of the phases (**Equation 2**). We compared this to the local order Kuramoto parameter (**Equation 4**) computed in the Schaefer1000 parcellation and compared to brain spirals in this space (obtained by averaging the corresponding values of the vertices within each parcel).

**Figure 3.**
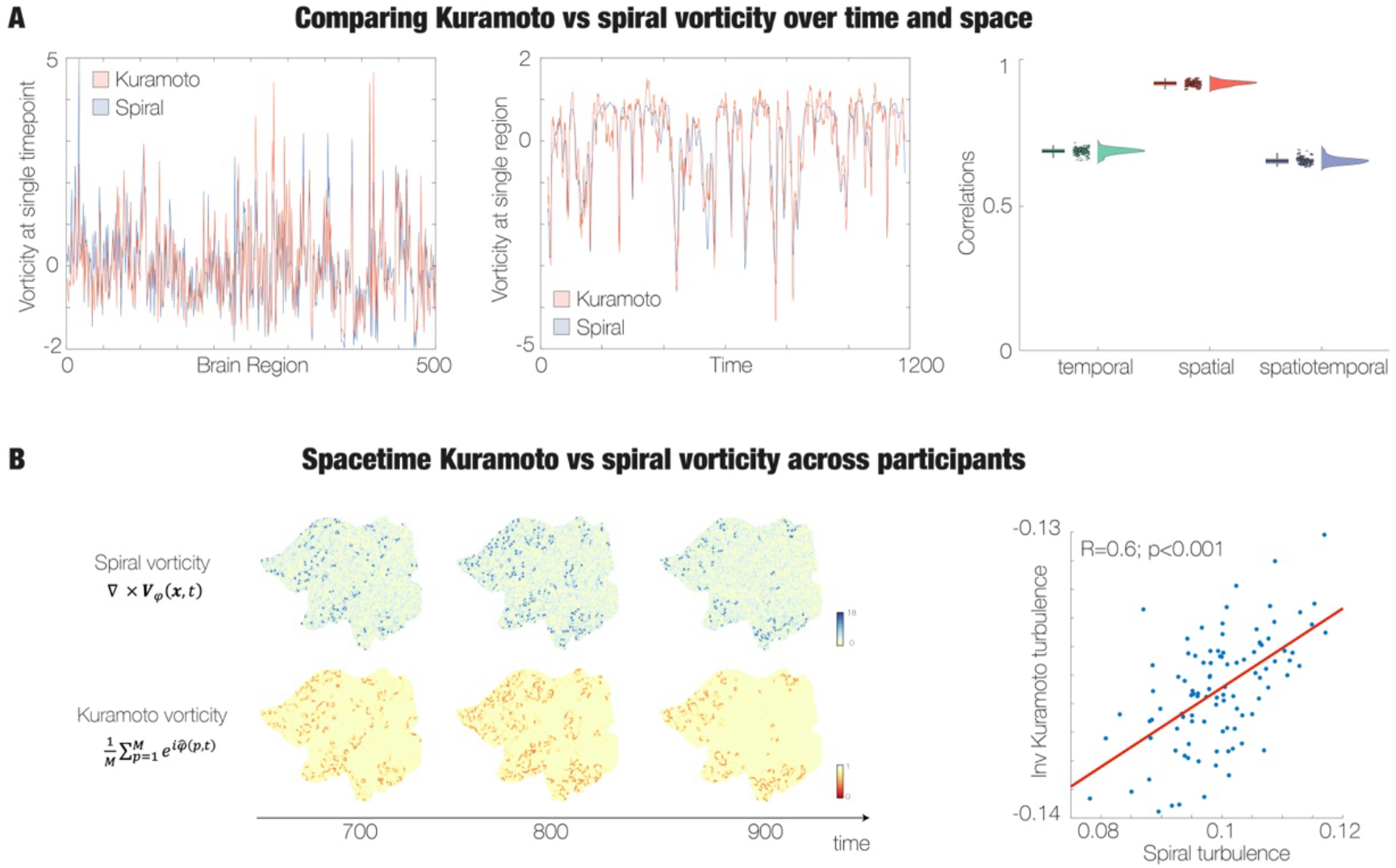
Equivalence between measures of Kuramoto and spiral vorticity. **A)** The left graph is comparing the measures of Kuramoto (red) and spiral (blue) vorticity. In order to show the similarity of the two measures, we z-scored the average of the values at a single time-point in the vertices within each of the parcels of the Schaefer1000 parcellation. Please note that Kuramoto values are inverted to improve the visualisation of the similarities. Here, the graph only shows these values for one hemisphere. Similarly, the middle panel is showing the z-scored values of a single parcel over the full time (1200 TRs). The right panel is showing a boxplot of the correlations between Kuramoto and spiral vorticity across time (green), space (red) and spacetime (blue) for a subset of 100 HCP participants. Please note the generally high correlations across all three, with the highest correlation found in the spatial domain. **B)** The left panel visualises the equivalence between spiral and Kuramoto vorticity by showing three snapshots of the vorticity in the full vertex space of an individual. This high spatiotemporal correlation can also be observed across participants, using the measures of spiral and Kuramoto turbulence. The graph shows the high correlation between both measures of turbulence.

**Figure 3A** (left panel) shows the results of comparing the measures of Kuramoto (red) and spiral (blue) vorticity across one hemisphere of 500 parcels. Note that the two measures are inverted in terms of measuring synchronisation, that is the lowest level of synchronisation is found at the centre of the brain spiral, while the opposite is true for Kuramoto vorticity (compare **Equations 2** and **4**). Therefore, in the graph we have inverted the Kuramoto vorticity values and z-scored both values to improve the visualisation of their similarities. In the middle graph, the figure shows the similarity in terms of time by showing the z-scored values of a single parcel over the full time (1200 TRs). The right panel shows boxplots as quantification of the correlations between the two measures in terms of time (green), space (red) and spacetime (blue) for a subset of 100 HCP participants. The highest correlation is found in the spatial domain.

**Figure 3B** shows these spatial similarities very clearly through three snapshots of the two kinds of vorticity (spiral vorticity in green and Kuramoto vorticity in yellow) in the full vertex space in an individual. In order to further quantify these similarities, we also computed the spiral turbulence (**Equation 3**) and the Kuramoto turbulence (**Equation 5**). As can be seen from the scatterplot in the right panel, this high spatiotemporal correlation is also found across the 100 participants (R=0.57, p<0.001).

### Whole-brain modelling using partition

Having shown the equivalence between spiral and Kuramoto vorticity, we proceed to model interactions between the Kuramoto vortices. In order to do so, we simplified the problem through using a common method for space partitioning (Snyder *et al*., 2020), creating a lower dimensional vortex space that can be used for modelling. We demonstrated the effectiveness of this approach from first principles by deriving the whole-brain model in Schaefer100 vortex space from a phase model in the finer Schaefer1000 partition (see *Methods*).

This partitioning is schematised in **Figure 4**. First, we show renderings of the Schaefer1000 parcellation on a flattened hemisphere (top) as well as the phases of BOLD signals giving rise to the FC phaselock matrix (**Figure 4A**). This is used in a whole-brain model of the FC phaselock matrix using the Kuramoto phase model for the local dynamics (Cabral *et al*., 2014) (**Figure 4B**). Using the partitioning method we can derive a closed equation for the vortices in the coarser Schaefer100 vortex space. **Figure 4C** shows renderings of the Schaefer100 parcellation in a flattened hemisphere (top) and with the values of Kuramoto vorticity in this parcellation (bottom left), giving rise the FC vorticity matrix (bottom right). As shown in **Figure 4D**, this is then used in a whole-brain model derived analytically from finer parcellation Kuramoto model to produce a Hopf model in this coarser parcellation of the vortex space.

**Figure 4.**
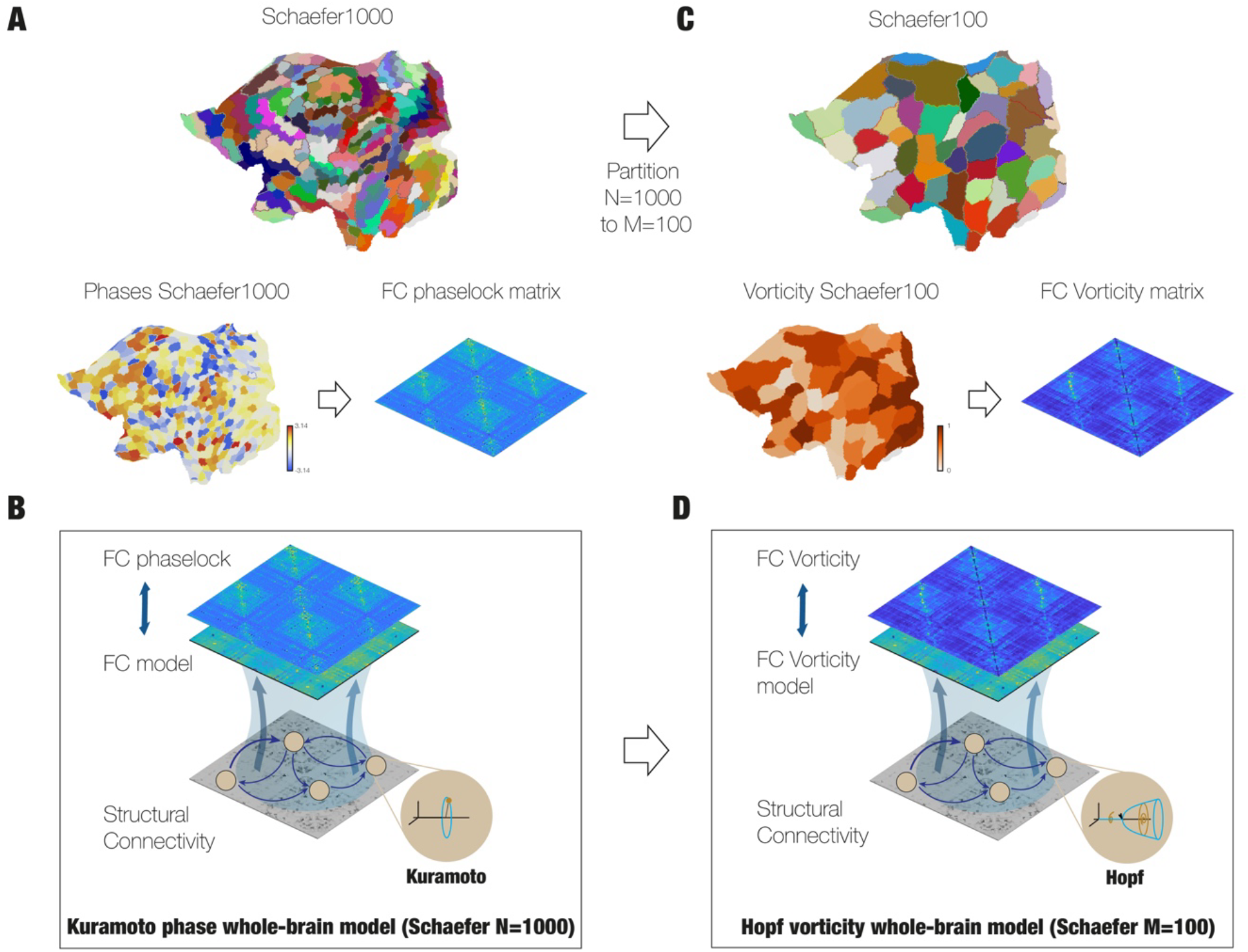
Whole-brain model in vortex space derived through space partitioning. A whole-brain model in vortex space can be derived from first principles from a phase model in a finer partition. **A)** First, for the whole-brain model in phase space, we use the fine Schaefer1000 partition (upper left part) to extract the corresponding phase lock matrix from the averaged signals from the empirical data in vertex space. **B)** A whole-brain model was built using a Kuramoto model for the local dynamics and coupled through the structural connectivity matrix for fitting the empirical phase lock matrix. **C)** Importantly, this model with Kuramoto local dynamics for the phases directly leads to an analytical whole-brain model with Stuart-Landau local dynamics of the vortex space (see Methods). Specifically, we define this Schaefer100 parcellation (top) as a partition of the Schaefer1000, where each parcel contains the Kuramoto vorticity (bottom left). The functional connectivity of this Kuramoto vorticity can then be used to fit the vortex whole-brain model. **D)** As shown the vortex whole-brain model is using the analytical version of the Hopf equation, now derived from first principles.

### Comparing Hopf vortex model vs Kuramoto phase model

**Figure 5** shows a direct comparison of the two models in vortex space. We measured this in terms of the fit of both models for the error (top) and structural similarity index measure (SSIM, bottom) between the generated and empirical functional connectivity in vortex space. To directly compare the two measures, we transformed the Kuramoto model from signal space to vortex space through a two-step process: 1) Optimise the Kuramoto phase model to the empirical phase lock data and 2) use this optimised Kuramoto phase model to generate time series and compute the corresponding functional connectivity in vortex space (see *Methods*).

**Figure 5.**
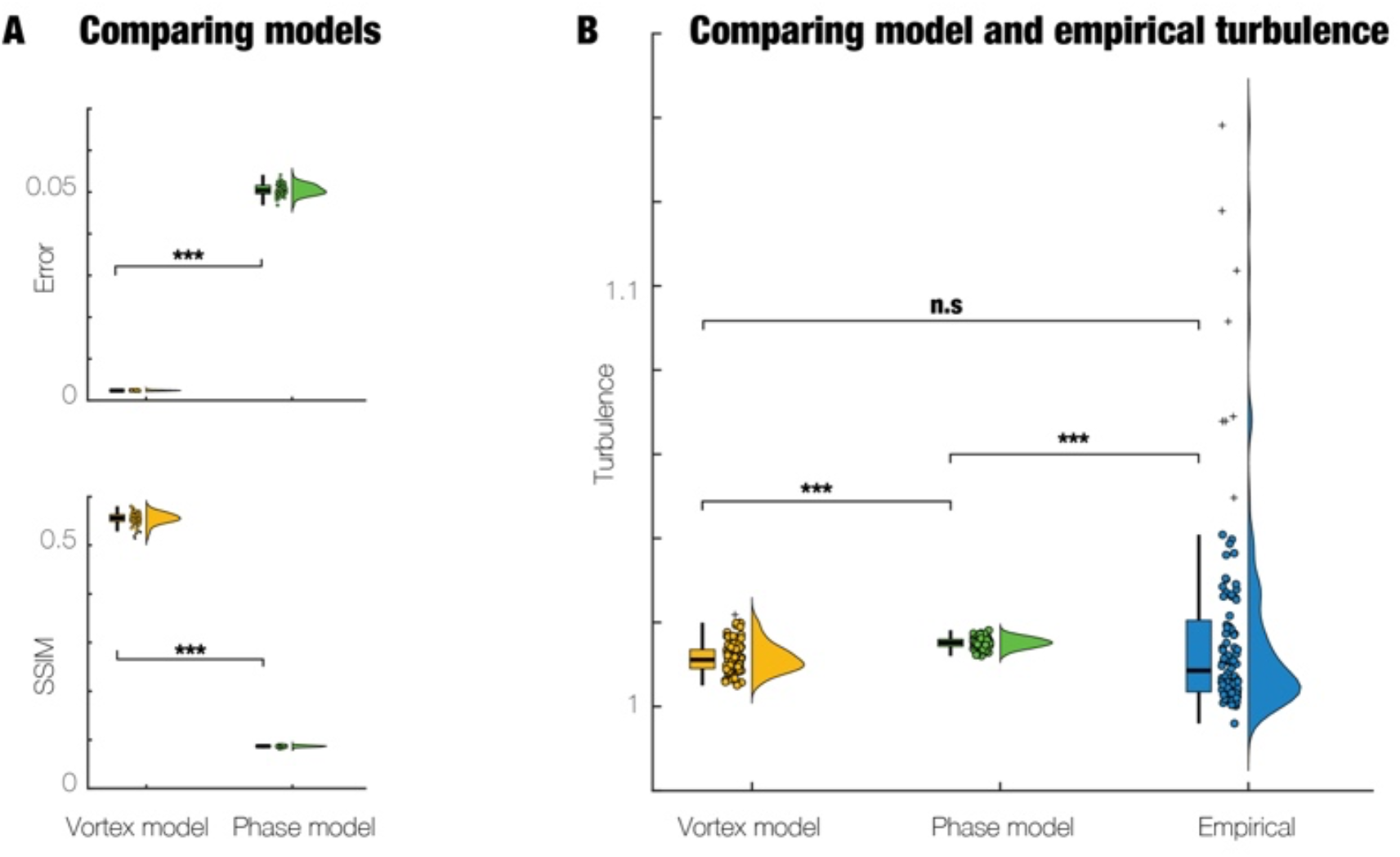
Hopf vortex whole-brain model is better than Kuramoto phase model for fitting empirical data. We compared the fit of both models in terms of the error (top) and Structural similarity index measure (SSIM, bottom) between the generated and empirical functional connectivity in vortex space. The Hopf vortex model is already in vortex space but the Kuramoto model is in signal space. So, in order to directly compare the accuracy of the two models, the Kuramoto model must be in vortex space. This is achieved by a two-step process: 1) Optimise the Kuramoto phase model to the empirical phase lock data and 2) use this optimised Kuramoto phase model to generate time series and compute the corresponding functional connectivity in vortex space (see Methods). **A)** The figure clearly shows that the Hopf vortex model significantly outperforms the Kuramoto phase model. **B)** Similarly, the Hopf vortex model was significantly better than the Kuramoto phase model in fitting the level of empirical turbulence. Importantly, there were no significant differences between the vortex model and the empirical data, which was not the case for the phase model.

**Figure 5A** shows that the Hopf vortex model significantly outperforms the Kuramoto phase model for the error (Wilcoxon *p<0*.*001*) and SSIM (Wilcoxon *p<0*.*001*). Similarly, **Figure 5B** shows that the Hopf vortex model was significantly better than the Kuramoto phase model for fitting the level of empirical turbulence (Wilcoxon *p<0*.*001*). In fact, there were no significant differences between the turbulence of the data generated by the vortex model and turbulence of the empirical data. In contrast this was not true for the phase model, demonstrating the superiority of the vortex model over the phase model.

### Cognition can be described by whole-brain model of the interactions of vortices

As mentioned in the introduction, many attempts have been made to find adequate measures for successfully decoding and distinguishing significant aspects human cognitive processing. Here we were able to show that whole-brain models of the interactions between vortices can be very useful for classification and understanding the mechanisms for the underlying computation.

**Figure 6** shows the results of using the Hopf vortex whole-brain model to quantify the vortex interactions. In particular, the model provides an optimised individual measure of the generative effective connectivity (GEC) matrix, which is a direct measure of the interactions between vortices (see *Methods*). This is then used on two different conditions of resting state and SOCIAL task. For this we used SVM classification with the individual GECs for both conditions for all 971 HCP participants. This showed a near perfect classification on the generalisation set with 0.975±0.025 (mean±s.d.) accuracy on the 1,000-folds. This is also shown in the confusion matrix and in the boxplot across the folds (**Figure 6A**). The differences in regional degree of the average GEC for rest and SOCIAL task are shown in brain renderings (**Figure 6B**).

**Figure 6.**
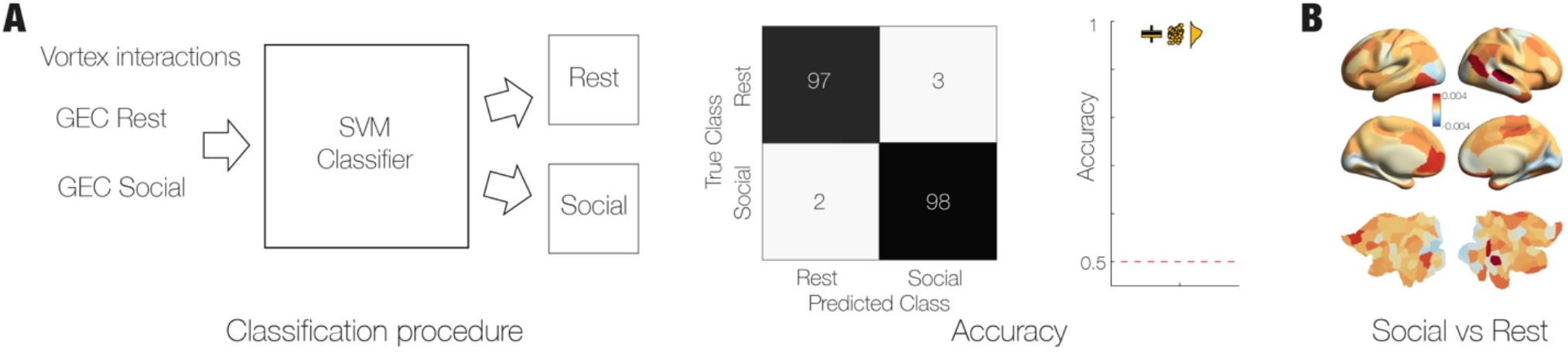
Classification of vortex interactions is highly efficient for distinguishing rest vs social task. Using the Hopf vortex whole-brain model can be used to quantify the vortex interactions in the form of the generative effective connectivity (GEC) for different conditions (rest and SOCIAL task). **A)** We classified the individual GECs corresponding to rest and social for all 971 participants and found a near perfect classification on the generalisation set with 0.975±0.025 (mean±s.d.) accuracy on the 1,000-folds. This is shown in the confusion matrix and in the boxplot across the folds. **B)** Rendering of the differences in regional degree of the average GEC for rest and SOCIAL task.

Importantly, this whole-brain modelling is not just useful for distinguishing tasks from rest but is also highly accurate in classifying the subtleties between sub-tasks. **Figure 7** shows three examples of using the Hopf vortex whole-brain model to quantify the vortex interactions in the form of the generative effective connectivity (GEC) for different subtle sub-task conditions.

**Figure 7.**
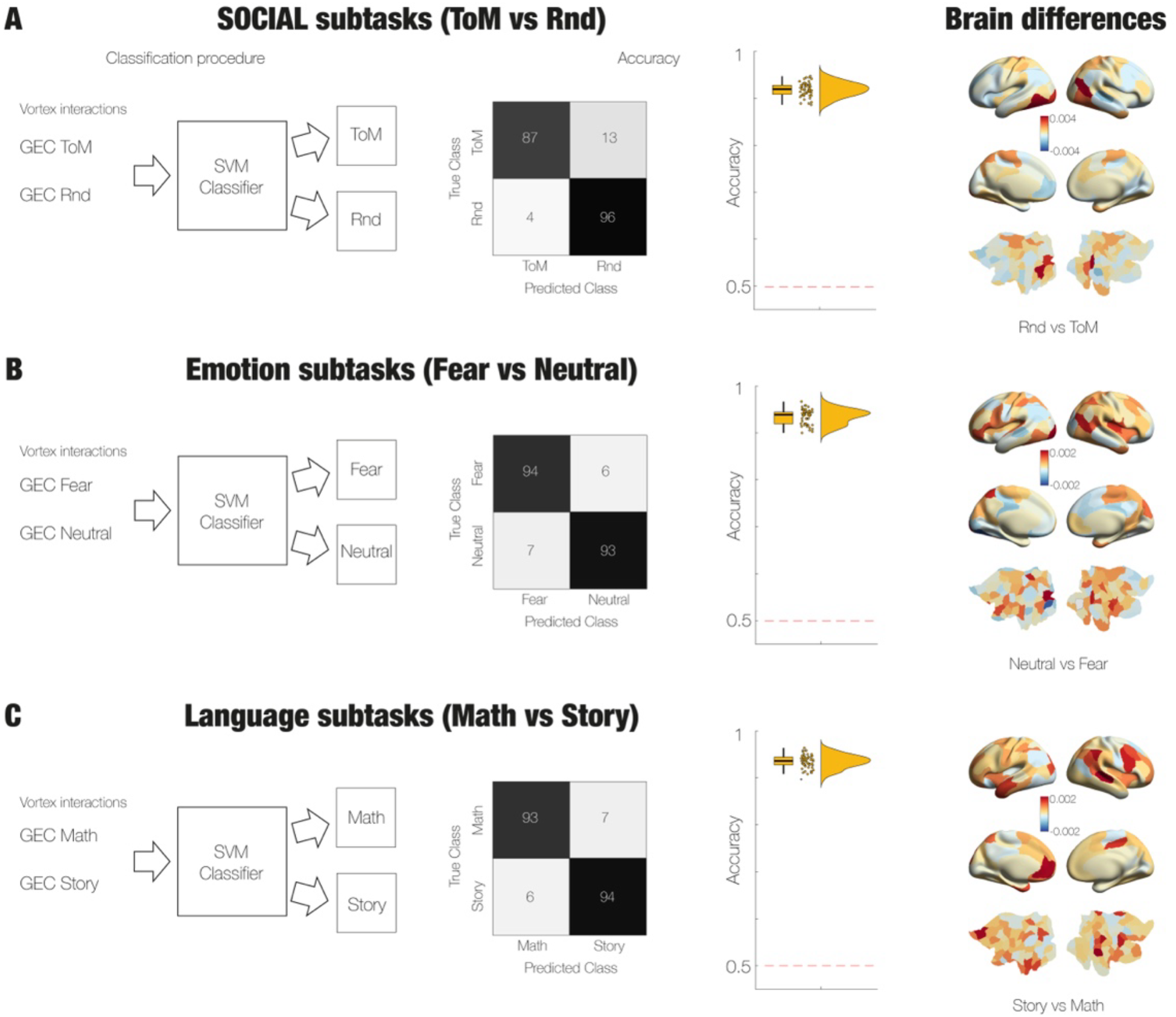
Interactions in turbulent vortex space are highly accurate in classifying subtle sub-tasks. Similar to figure 6, we used the Hopf vortex whole-brain model to quantify the vortex interactions in the form of the generative effective connectivity (GEC) for different subtle sub-task conditions. **A)** Within the SOCIAL task, which uses moving dots on a screen, there are two conditions, namely moving dots evoking theory of mind (ToM) and randomly moving dots (Rnd). We found a near perfect classification on the generalisation set with 0.92±0.02 (mean±s.d.) accuracy. The rendering shows the differences in regional degree of the average GEC for ToM vs Rnd sub-tasks. **B)** Same but for the Fear and Neutral sub-tasks of the EMOTION task. We found a near perfect classification on the generalisation set with 0.94±0.02 (mean±s.d.) accuracy. The rendering shows the differences in regional degree of the average GEC for Fear vs Neutral sub-tasks. **C)** Similar but for the Maths and Story sub-tasks of the LANGUAGE task. We found a near perfect classification on the generalisation set with 0.94±0.02 (mean±s.d.) accuracy. The rendering shows the differences in regional degree of the average GEC for Maths vs Story sub-tasks.

**Figure 7A** shows the near perfect classification on the generalisation set with 0.92±0.02 (mean±s.d.) accuracy of comparing the two conditions in the SOCIAL task. This tasks uses moving dots on a screen with two distinct conditions of the moving dots either evoking theory of mind (ToM) or moving randomly (Rnd). The brain renderings show the differences in regional degree of the average GEC for ToM vs Rnd sub-tasks.

**Figure 7B** shows the results of doing the same comparison for the Fear and Neutral sub-tasks of the EMOTION task. Again, a near perfect classification was found on the generalisation set with 0.94±0.02 (mean±s.d.) accuracy. The brain rendering shows the differences in regional degree of the average GEC for Fear vs Neutral sub-tasks.

Finally, **Figure 7C** shows the results of classifying the Maths and Story sub-tasks of the LANGUAGE task. A near perfect classification was found on the generalisation set with 0.94±0.02 (mean±s.d.) accuracy. The brain rendering shows the differences in regional degree of the average GEC for Maths vs Story sub-tasks.

These results leave open the intriguing possibility of controlling the turbulent brain dynamics in vortex space, similar to how fluid dynamics can be efficiently controlled in vortex space. In a proof of principle, we were able to find model perturbations in vortex space that led to a transitioning between resting and the SOCIAL task. Similar to the perturbation strategy we have previously used in signal space (Deco *et al*., 2019), here we used noise perturbations of the local dynamics of the Hopf model in vortex space to force a transition between resting and SOCIAL task. We were able to find a set of perturbation that transformed the dynamics of the model resting state to the SOCIAL task with a model fit correlation of 0.67 to the empirical FC of the SOCIAL task.

## Discussion

Here we have shown that whole-brain modelling of the interactions of turbulent vortices provides an excellent framework for quantifying and understanding brain dynamics. This is important since turbulence is a fundamental principle found at all scales in Nature, from the microscopic scale to the scale of the universe. The recent discovery of turbulence in brain dynamics provides a mechanistic basis of highly efficient time-critical information processing and transmission across wide-spread brain networks. Here we have provided the first evidence that the interactions between turbulent vortices could be the fundamental principle of brain computation.

We quantified these interactions through building a whole-brain model of the vortex space and found that it not only successfully distinguishes the detailed spacetime dynamics of rest and cognition, but that it can even distinguish between subtle sub-components of cognitive tasks. Importantly, this Hopf whole-brain model in vortex space was derived from first principles from the fine-grained vertex space of interacting coupled Kuramoto phase oscillators. Overall, this provides a natural vortex space for brain computation and could potentially offer new ways of controlling the breakdown in disease.

The whole-brain modelling in vortex space was made possible by providing a mapping between the signal space in a fine-grained space of 32k vertices to a coarser-grained vortex space of 100 regions. For the necessary partitioning scheme, we used the well-known method by Snyder and colleagues (Snyder *et al*., 2020) which made it possible to derive analytically the whole-brain model of the vortex space from using a model with the Kuramoto phase oscillator in the vertex space. We were able to show that this results in a model using the Hopf (or Stuart-Landau) oscillator for the local dynamics in vortex space.

This result can be seen in the wider context of the use of Hopf whole-brain models, which, set at the edge of the bifurcation point and coupled to anatomical connectivity of the connectome, have been shown to be able to model the signal space of brain dynamics (Deco *et al*., 2017). The Hopf model is highly efficient for capturing the stochastically fluctuating signal with oscillatory components found in the slow dynamics of BOLD signals. The reason for this ability can be found in the fact that a whole-brain model using local dynamics of the exact mean-field model with inhibitory and excitatory integrate-and-fire neurons lead to similar dynamics as the Hopf model, poised just below the critical Hopf bifurcation (Perl *et al*., 2023b). Furthermore, Piccinini and colleagues found that from all possible connectome-based models, human fMRI was best described by a Hopf model just below the bifurcation point (Piccinini *et al*., 2022). This matches the conclusions of Sip and colleagues (Sip *et al*.) who found that the regional and network-level dynamics of resting-state fMRI can be described as noisy fluctuation around a single fixed point, which again is best captured by the Hopf model. As such, this body of research has convincingly shown the Hopf model is superior to what is often termed as more biologically detailed models of excitation and inhibition.

Still, the physics of fluid turbulence has shown that the best models are not based at the signal level but rather at the higher vortex level. The rotationality of the vortices provide sufficient information to understand and ultimately control turbulence. Our results provide a similar reason to move beyond the signal level for the oscillatory turbulence found in brain dynamics. The partitioning method used here clearly demonstrates that the Hopf whole-brain model emerges as the natural framework for modelling brain turbulence. In particular, using the vortex space for the whole-brain model provides an individualised an explicit description of the interactions of the brain vorticity through the generative effective connectivity (GEC).

The results here show the utility of this individualised GEC since it is excellent not only for classification of rest versus SOCIAL task but also for distinguishing between very similar sub-phases within tasks. However, this is not just useful for classification but also provides the necessary elements for a mechanistic region-level explanation of task computation. Take for example the differences between the interactions during resting state and performing the SOCIAL task of assigning meaning to dots moving around a computer screen and deciding whether these reflect some degree of theory of mind or simply random movement. Here there are distinct differences in the brain networks engaged, with clear differences in the task related activity in visual regions and in the medial prefrontal cortex for rest and SOCIAL task.

Overall, modelling the interactions between turbulent vortices is a powerful way of understanding brain dynamics in health. But equally, this also opens the possibility of controlling brain dynamics in disease. Excitingly, in a proof of principle, we were able to perturb the whole-brain model of resting state to instead fit the SOCIAL task condition with a high level of precision. This opens up for a much needed ability to control turbulence in disease. Future studies could investigate data from patients with for example traumatic brain injury, PTSD and depression to design therapeutic interventions to stop full-blown attack with potential catastrophic impact on the whole-brain dynamics (Figueroa *et al*., 2019; Hinojosa *et al*., 2024; Jirsa *et al*., 2023; Martinez-Molina *et al*., 2024) by changing the interactions between vortices to a healthy regime. Longer-term, this could have important impact on providing new therapies for current treatment-resistant neuropsychiatric disorders.

## Methods

### Empirical fMRI data

#### Neuroimaging Ethics HCP

The Washington University–University of Minnesota (WU-Minn HCP) Consortium obtained full informed consent from all participants, and research procedures and ethical guidelines were followed in accordance with Washington University institutional review board approval (Mapping the Human Connectome: Structure, Function, and Heritability; IRB # 201204036)..

#### Neuroimaging HCP Participants

The data set used for this investigation was selected from the March 2017 public data release from the Human Connectome Project (HCP) where we chose a sample of 971 participants with data from resting state and all seven cognitive tasks.

#### Neuroimaging acquisition for fMRI HCP

The 971 HCP participants were scanned on a 3-T connectome-Skyra scanner (Siemens). We used one resting state fMRI acquisition of approximately 15 minutes acquired on the same day, with eyes open with relaxed fixation on a projected bright cross-hair on a dark background as well as data from the seven tasks. The HCP website (http://www.humanconnectome.org/) provides the full details of participants, the acquisition protocol and preprocessing of the data for both resting state and the seven tasks.

#### Preprocessing and extraction of functional timeseries in fMRI resting data

The preprocessing of the HCP resting state and task datasets is described in details on the HCP website. Briefly, the data is preprocessed using the HCP pipeline which is using standardised methods using FSL (FMRIB Software Library), FreeSurfer, and the Connectome Workbench software (Glasser *et al*., 2013; Smith *et al*., 2013). This preprocessing included correction for spatial and gradient distortions and head motion, intensity normalization and bias field removal, registration to the T1 weighted structural image, transformation to the 2mm Montreal Neurological Institute (MNI) space, and using the FIX artefact removal procedure (Navarro Schroder *et al*., 2015; Smith *et al*., 2013). The head motion parameters were regressed out and structured artefacts were removed by ICA+FIX processing (Independent Component Analysis followed by FMRIB’s ICA-based X-noiseifier) (Griffanti *et al*., 2014; Salimi-Khorshidi *et al*., 2014). Preprocessed timeseries of all grayordinates are in HCP CIFTI grayordinates standard space and available in the surface-based CIFTI file for each participants for resting state and each of the seven tasks.

We used a custom-made Matlab script using the ft_read_cifti function (Fieldtrip toolbox (Oostenveld *et al*., 2011)) to extract the average timeseries of all the grayordinates in each region of the Schaefer parcellation, which are defined in the HCP CIFTI grayordinates standard space. Furthermore, the BOLD time series were transformed to phase space by filtering the signals in the range between 0.008-0.08 Hz, where we chose the typical highpass cutoff to filter low-frequency signal drifts (Fox *et al*., 2005), and the lowpass cutoff to filter the physiological noise, which tends to dominate the higher frequencies. We then applied the Hilbert transforms in order to obtain the phases of the signal for each brain node as a function of the time.

### The HCP task battery of seven tasks

The HCP task battery consists of seven tasks: working memory, motor, gambling, language, social, emotional, relational, which are described in details on the HCP website (Barch *et al*., 2013). HCP participants performed all tasks in two separate sessions (first session: working memory, gambling and motor; second session: language, social cognition, relational processing and emotion processing).

#### Schaefer parcellations

Schaefer and colleagues created a publicly available population atlas of cerebral cortical parcellation based on estimation from a large data set (N = 1489) (Schaefer *et al*., 2018). They provide parcellations of 100 and 1000 areas available in surface spaces, as well as MNI152 volumetric space. We used here the Schaefer parcellations with 1000 and 100 parcels and extracted the timeseries from HCP using the HCP surface space version.

### Structural connectivity using dMRI

The Human Connectome Project (HCP) database contains diffusion spectrum and T2-weighted imaging data from 32 participants with the acquisition parameters described in detail on the HCP website (Setsompop *et al*., 2013). The freely available Lead-DBS software package (http://www.lead-dbs.org/) provides the preprocessing which is described in detail in Horn and colleagues (Horn *et al*., 2017) but briefly, the data was processed using a generalized q-sampling imaging algorithm implemented in DSI studio (http://dsi-studio.labsolver.org). Segmentation of the T2-weighted anatomical images produced a white-matter mask and co-registering the images to the b0 image of the diffusion data using SPM12. In each HCP participant, 200,000 fibres were sampled within the white-matter mask. Fibres were transformed into MNI space using Lead-DBS (Horn and Blankenburg, 2016). We used the standardised methods in Lead-DBS to produce the structural connectomes for both Schaefer1000 and Schaefer100 parcellation schemes.

### Theoretical turbulent vortices framework

#### Spiral vorticity

Generating the spiral vortices requires computing the phase map of the empirical fMRI signals in the 2D flattened cortical space and then calculating the phase vector field, i.e. the phase velocity flow, whose curl defines the spirals vorticity. These spirals characterise local rotational motion near a specific point, i.e. the tendency to rotate, as observed from above at that point and moving with the flow. For a subset of 100 participants of the HCP database described above, we consider 2 mm standard CIFTI grayordinate space of half hemisphere comprising 32k cortical vertices (vertex space) which are represented in the 2D flattened cortex surface. Temporal filtering of the BOLD fMRI signals is done by applying the second-order zero-phase Butterworth filter (0.008 Hz < f < 0.08 Hz) to each voxel of the cortex to focus on slow neuronal fluctuations. After this, the data is smoothed by performing a Gaussian spatial filtering with a width of 4 mm. Finally, for each time point *t*, we compute the phase map *φ*(***x***, *t*) in the 2D vertex space ***x*** using the Hilbert transform. The phase vector field is computed as the spatial gradient of the phase map given by:

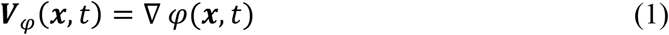

The spatial gradient was determined by taking derivatives across the two spatial dimensions, numerically computed by using central finite differences. Circular statistics were used to compute the differences. Finally, we derived the spiral vorticity in vertex space by computing the spatial curl of the phase vector field, that is:

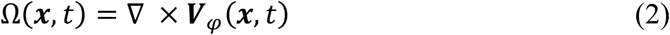

The curl is thresholded at Ω >1 or Ω <−1 to find the potential core of spirals with very high vorticity. We define spiral based turbulence *T*_*spiral*_ as the standard deviation of the spiral vorticity map across space and time:

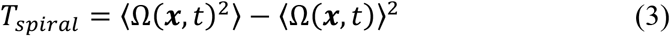

where the brackets ⟨ ⟩ denotes average across vertex space and time.

#### Kuramoto vorticity

As has been shown, turbulence is not restricted to fluid dynamics but is also found in other physical systems including coupled oscillators (Kawamura *et al*., 2007) and brains (Deco and Kringelbach, 2020; Deco *et al*., 2023; Deco *et al*., 2025). Yoshiki Kuramoto used the theory of coupled oscillators to show turbulence in fluid dynamics (Kuramoto, 1984), suggestive of how turbulence could be important not only for energy transfer but for efficient information transfer. Specifically, this framework defines the Kuramoto local order parameter, representing a spatial average of the complex phase factor of the local oscillators weighted by the coupling. Thus, the Kuramoto local order parameter gives the level of synchronisation of the local phases around a specific point. This is we here call *Kuramoto vorticity*. Importantly, the level of amplitude turbulence is defined as the standard deviation of the modulus of Kuramoto vorticity and can be applied to the empirical data of any physical system. Remarkably, brains also were also found to exhibit a similar turbulent power law, strongly suggesting the presence of a cascade of efficient information processing across scales (Deco and Kringelbach, 2020; Deco *et al*., 2023; Deco *et al*., 2025).

As can be appreciated from **Figure 2**, both spirals and Kuramoto vorticity show the same phenomenon, namely the local rotation of the moving flow, albeit in different complementary ways. Specifically, the Kuramoto vorticity in the 2D vertex space, *R*(***x***, *t*), is defined by:

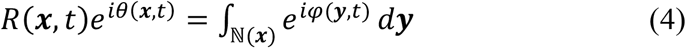

where *φ*(***y***, *t*) is the local phase of the BOLD signal in the 2D vertex space at point ***y***. The integral range ℕ(***x***) is defined by a circle of radius 4 mm around ***x***. We measure amplitude turbulence by first defining the Kuramoto local order parameter and then taking the standard deviation of the modulus across time and space (Kawamura *et al*., 2007). Thus, the Kuramoto turbulence, *T*_*Kuramoto*_, is defined as the standard deviation of the Kuramoto vorticity across space and time:

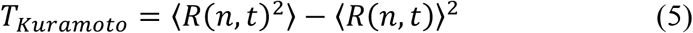

where the brackets ⟨ ⟩ denotes average across vertex space and time.

In order to directly compare Spiral vorticity and Kuramoto vorticity, we move from vertex space to the more coarse-grained Schaefer1000 parcellation by averaging the values of the vertices within each parcel in the subset of 100 HCP participants used for the spiral vorticity.

### Kuramoto vorticity whole-brain model

A key finding of the coarse-grained partition methodology is the derivation of a model reduction of a coupled system of Kuramoto oscillators (Snyder *et al*., 2020). One of the most well-known examples of this kind of model reduction is the groundbreaking work by Ott and Antonsen (Ott and Antonsen, 2008), who demonstrated that in the limit of an infinite number of oscillators, the overall synchronisation level follows an autonomous ordinary differential equation. In neuroscience, research has robustly demonstrated that a system of Kuramoto oscillators coupled through the structural anatomical connectivity matrix is able to achieve an excellent fitting of functional brain activity obtained with MEG (Cabral *et al*., 2014).

Given that the vortices are fundamental elements of the brain dynamics associated with the processing and computation of cognition, it would be convenient to derive a direct reduced model of the vortical activity emerging from an underlying Kuramoto whole-brain model. Furthermore, if the underlying base Kuramoto whole-brain model is defined in a fine parcellation (in our case the Schaefer1000), the reduced model of the vortical activity will conveniently be defined in a more coarse parcellation (here Schaefer100).

Let us consider a Kuramoto whole-brain model in the Schaefer1000 parcellation. The models consist of a network of *M* = 1000 coupled phase oscillators, where the connections are defined by the anatomical connectivity matrix, which was estimated using DMRI data. This model assumes that oscillators interact based on their phase differences. Let 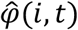 represent the phase of the *i-th* oscillator at time *t*. We will use the circumflex hat to denote all the variables related to the Schaefer1000 parcellation with respect to the variables in vertex space (with circumflex). The phases evolve according to the following set of coupled differential equations:

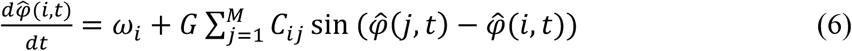

where *ω*_*i*_ is the natural frequency of the *i-th* oscillator, while *C*_*ij*_ is the anatomical structural connectivity matrix obtained by dMRI in the Schaefer1000 parcellation (see above), and *G* represents the global coupling strength. The natural frequency of oscillations for each ROI was estimated from the peak of the power spectra estimated from their BOLD in the frequency band 0.008–0.08 Hz. The interaction between two oscillators, *i* and *j*, is governed by the sine of their phase difference. This interaction promotes synchronisation, since an oscillator lagging behind (that is 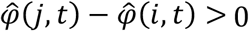) speeds up, while one ahead (that is 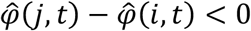) slows down. This model was numerically integrated using Euler’s method with a time step of 0.01 (equivalent to 10 ms).

One interesting order parameter of the system is the global Kuramoto order parameter, 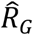, given by:

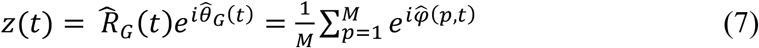

where 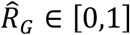 is the global synchrony and 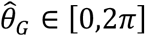 is the average phase. If all phases are equal then 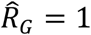, and if the phases are spread uniformly over the unit circle, then 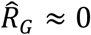. Thus 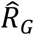 is a natural measure of synchronisation. It is interesting to note that in the case of *C*_*ij*_ = 1/*M*, the global Kuramoto order parameter follows a closed equation. If the frequencies *ω*_*i*_, there is, as demonstrated by Ott and Antonsen (Ott and Antonsen, 2008). Assuming that *ω*_*i*_ is Cauchy distributed, i.e., *ω*_*i*_~*f*(*ω*) where *f* is the Cauchy probability density function with mode Ω and width *δ*, and that the coupling is mean field, then 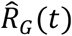 evolves according to a Stuart-Landau equation,

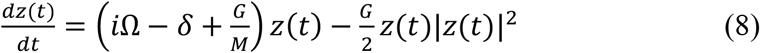

The last equation shows that *z* = 0 is always a solution but undergoes a pitchfork bifurcation at *G*_*critical*_ = 2*δ*, when a new solution with 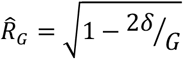, and 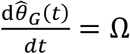 appears, representing partial synchrony that becomes global synchrony (i.e., 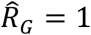) as *G* → ∞.

### Partitioning: Whole-brain model in vortex space

Importantly, a very similar analytical derivation can be carried out for the case where oscillators are not all coupled to each other but are instead grouped into subsets, with the coupling strength between any two oscillators depending on the subsets they belong to. In other words, a close equation can be found for the local Kuramoto order parameters evolution, that is for the dynamics of the vortex space, defined in a partition of the original system. Here, the original system of Kuramoto oscillators was defined above in the Schaefer parcellation *M* = 1000.

We define a partition of that parcellation by adopting for the local Kuramoto order parameters a Schaefer parcellation *N* = 100. Let ℘ = (ℙ(1), …., ℙ(*N*)) represent a partition of the index set [1, …, *M*] and let *K* be a *N* × *N* matrix of coupling strengths. Following Snyder and colleagues (Snyder *et al*., 2020), such a modular system can be expressed as:

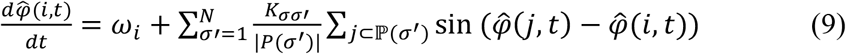

The same mathematical framework can be used to show that if, for each *σ*, the natural frequencies {*ω*_*i*_|*i* ⊂ ℙ(*σ*) are distributed according to a Cauchy distribution with mode Ω_*σ*_ and width *δ*_*σ*_, then the cluster order parameters *z*_*σ*_(*t*), defined by

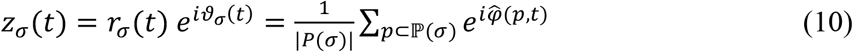

satisfy a coupled Stuart-Landau equation of the form:

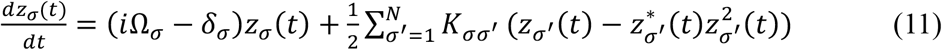

In this context, the natural emergence of last equation suggests that in the Kuramoto model, groups of oscillators can collectively behave as a single oscillator. This implies that there exists a renormalisation process for coupled oscillator systems that keeps them within the same model framework. Note, that the cluster order parameters *z*_*σ*_(*t*) are the Kuramoto vorticity on the Schaefer1000 parcellation defined by averaging spatially the complex phase factor of the local phases 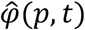 from the BOLD time series signals defined in this parcellation. The Kuramoto vorticity, *z*_*σ*_(*t*) reflect the level of synchronization of the local phases in a given partition *σ* (in the Schaefer100 parcellation). **Equation 11** is thus an explicit model of the vortex space.

Finally, in order to fit the vortex model to the empirical vortex data, we use for the simulations of the vortex space in the Schaefer100 parcellation following equation:

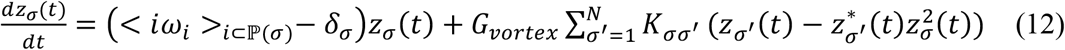

where *G*_*vortex*_ is a free parameter expressing the global coupling in vortex space, and 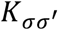 is the anatomical structural connectivity matrix obtained by dMRI in the Schaefer100 parcellation (see above). In the simulations we used *δ*_*σ*_ = 0.001 and added Gaussian noise with and standard deviation of 0.01. The diagonal elements of *K*_*σσ*_ were set to 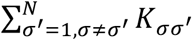 in order to recover the usual form of the Hopf whole-brain model (Deco *et al*., 2017), which provides a good level of fitting as shown in the results.

When we optimise the Kuramoto model, we fit the free parameter *G* (global coupling in the Schaefer1000 parcelation and BOLD space) by fitting the PLV (Phase Locking Value of the signal in the Schafer 1000 parcellation, using our previously published methods (Ponce-Alvarez *et al*., 2015), i.e. 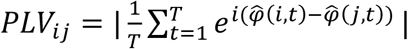 being *T* the maximal measured or simulated time, 1200 TR) and after finding the optimal *G* we compute the associated vortex space with **Equation 10** to compute the functional connectivity in vortex space as indicated below (**Equation 14**).

In order to directly compare the accuracy of the Kuramoto phase model with the Hopf vortex model, we compare the simulation functional connectivity in vortex space with the empirical functional connectivity vortex space.

### Capturing vortex interactions

In order to explicitly quantify the Kuramoto vortical interactions in the Schaefer100 parcellation we extend **Equation 12** to optimize instead the global conductivity parameter *G*_*vortex*_, the single paired conductivity parameters. The key idea is to fit the empirical functional connectivity of the Kuramoto vortices by the model described in **Equation 12**. This provides the Generative Effective Connectivity of the vortex space (GECVO) which is the effective weighting of the existing anatomical connectivity coupling the vortices in the Schaefer100 parcellation. Note that this is an extension of the classic concept of effective connectivity (Friston *et al*., 2003), where 1) GEC is generative, using the whole-brain model to adapt the strength of existing anatomical connectivity (i.e. the effective conductive values of each fibre) and 2) the optimisation target for GEC is in vortex space, that is the functional connectivity of the Kuramoto vortices. In other words, creating a whole-brain model in the vortex space of the empirical neuroimaging data provides direct access to determining the interactions between vortices.

We optimized GECVO between brain regions by comparing the output of the vortex model:

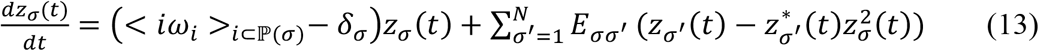

with the empirical measures of functional connectivity in vortex space. Let us denote by 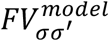 and 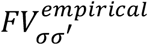 the simulated and empirical functional connectivity in vortex space, respectively. In our definition of functional connectivity in vortex space we considered the conjugated correlations as following:

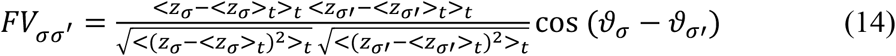

where the brackets ⟨ ⟩_*t*_ denotes average across time. In the model case the Kuramoto vortices are simulated by using **Equation 13**. In the empirical case the Kuramoto vortices are computed by using the Hilbert transform of the filtered empirical BOLD data (second-order zero-phase Butterworth filter: 0.008 Hz < f < 0.08 Hz) in the Schaefer1000 parcellation and defined as in **Equation 10**, but instead using the empirical extracted phases.

Using a heuristic gradient algorithm, we proceed to update the GEC such that the fit is optimised. In order to work only positive values for the algorithm, all values are transformed into a mutual information measure (assuming Gaussianity). More specifically, the updating uses the following form:

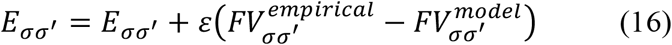

The model was run repeatedly with the updated GEC until the fit converges towards a stable value. In each iteration step, as before, the diagonal elements of *E*_*σσ*_ were set to 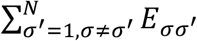.

We initialised using the anatomical connectivity (obtained with probabilistic tractography from dMRI) and only update known existing connections from this matrix (in either hemisphere). However, there is one exception to this rule which is that the algorithm also updates homologue connections between the same regions in either hemisphere, given that tractography is known to be less accurate when accounting for this connectivity. We used *ε* = 0.01, and continue until the algorithm converges. For each iteration we compute the model results as the average over as many simulations as there are participants. In the simulations we used *δ*_*σ*_ = 0.001 and added Gaussian noise with and standard deviation of 0.01.

### Support vector machine used for condition classification

We used a support vector machine (SVM) with polynomial kernels as implemented in the Matlab function *fitcecoc*. The function returns a fully trained, multiclass, error-correcting output codes (ECOC) model. This is achieved using the predictors in the input with class labels. The SVM used inputs of the 100×100 matrices of model GECVO, while the output was two classes corresponding to the conditions. We used the output from all 971 HCP participants used for generalisation, subdivided into 90% training and 10% validation, repeated and shuffled 1,000 times.

## Acknowledgments

G.D. is supported by Grant PID2022-136216NB-I00 funded by MICIU/AEI/10.13039/501100011033 and by “ERDF A way of making Europe”, ERDF, EU, Project NEurological MEchanismS of Injury, and Sleep-like cellular dynamics (NEMESIS) (ref. 101071900) funded by the EU ERC Synergy Horizon Europe, and AGAUR research support grant (ref. 2021 SGR 00917) funded by the Department of Research and Universities of the Generalitat of Catalunya. Y.S.P. is supported by was supported by the project NEurological MEchanismS of Injury, and Sleep-like cellular dynamics (NEMESIS) (ref. 101071900) funded by the EU ERC Synergy Horizon Europe. M.L.K. is supported by the Centre for Eudaimonia and Human Flourishing (funded by the Pettit and Carlsberg Foundations) and Center for Music in the Brain (funded by the Danish National Research Foundation, DNRF117). The funders had no role in study design, data collection and analysis, decision to publish or preparation of the manuscript.

## Code

The open-source MATLAB code can be found here and HCP data is in the public domain: https://github.com/decolab/TTB_modellingvortexinteraction

## References

Barch, D. M., Burgess, G. C., Harms, M. P., Petersen, S. E., Schlaggar, B. L., Corbetta, M., Glasser, M. F., Curtiss, S., Dixit, S., Feldt, C., Nolan, D., Bryant, E., Hartley, T., Footer, O., Bjork, J. M., Poldrack, R., Smith, S., Johansen-Berg, H., Snyder, A. Z., Van Essen, D. C. and Consortium, W. U.-M. H. (2013) Function in the human connectome: task-fMRI and individual differences in behavior. NeuroImage 80, 169–189.

Bewley, G. P., Lathrop, D. P. and Sreenivasan, K. R. (2006) Superfluid helium: visualization of quantized vortices. Nature 441, 588.

Cabral, J., Luckhoo, H., Woolrich, M., Joensson, M., Mohseni, H., Baker, A., Kringelbach, M. L. and Deco, G. (2014) Exploring mechanisms of spontaneous functional connectivity in MEG: How delayed network interactions lead to structured amplitude envelopes of band-pass filtered oscillations. Neuroimage 90, 423–435.

Christoph, J., Chebbok, M., Richter, C., Schroder-Schetelig, J., Bittihn, P., Stein, S., Uzelac, I., Fenton, F. H., Hasenfuss, G., Gilmour, R. F., Jr. and Luther, S. (2018) Electromechanical vortex filaments during cardiac fibrillation. Nature 555, 667–672.

Deco, G., Cruzat, J., Cabral, J., Tagliazucchi, E., Laufs, H., Logothetis, N. K. and Kringelbach, M. L. (2019) Awakening: predicting external stimulation forcing transitions between different brain states. PNAS 116, 18088–18097.

Deco, G. and Kringelbach, M. L. (2020) Turbulent-like dynamics in the human brain. Cell Reports 33, 108471.

Deco, G., Kringelbach, M. L., Jirsa, V. and Ritter, P. (2017) The Dynamics of Resting Fluctuations in the Brain: Metastability and its Dynamical Core [bioRxiv 065284]. Scientific Reports 7, 3095.

Deco, G., Leibana Garcia, S., Sanz Perl, Y., Tagliazucchi, E. and Kringelbach, M. L. (2023) The effect of turbulence in brain dynamics information transfer measured with magnetoencephalography. Commun Phys 6, 74.

Deco, G., Perl, Y. S., Jerotic, K., Escrichs, A. and Kringelbach, M. L. (2025) Turbulence as a framework for brain dynamics in health and disease. Neurosci Biobehav Rev 169, 105988.

Escrichs, A., Sanz Perl, Y., Fisher, P. M., Martinez-Molina, N. E. G. G., Frokjaer, V. G., Kringelbach, M. L., Knudsen, G. M. and Deco, G. (2024) Whole-brain turbulent dynamics predict responsiveness to pharmacological treatment in major depressive disorder. Mol Psychiatry 30, 1069–1079.

Escrichs, A., Sanz Perl, Y., Uribe, C., Camara, E., Turker, B., Pyatigorskaya, N., Lopez-Gonzalez, A., Pallavicini, C., Panda, R., Annen, J., Gosseries, O., Laureys, S., Naccache, L., Sitt, J. D., Laufs, H., Tagliazucchi, E., Kringelbach, M. L. and Deco, G. (2022) Unifying turbulent dynamics framework distinguishes different brain states. Commun. Biol. 5, 638.

Figueroa, C. A., Cabral, J., Mocking, R. J. T., Rapuano, K., van Hartevelt, T., Deco, G., Schene, A. H., Kringelbach, M. L. and Ruhe, H. G. (2019) Altered ability to access a clinically relevant control network in patients remitted from Major Depressive Disorder. HBM 10, 2771–2786.

Fox, M. D., Snyder, A. Z., Vincent, J. L., Corbetta, M., Van Essen, D. C. and Raichle, M. E. (2005) The human brain is intrinsically organized into dynamic, anticorrelated functional networks. Proc. Natl. Acad. Sci. U.S.A. 102, 9673–9678.

Frisch, U. (1995) Turbulence: The Legacy of A. N. Kolmogorov. Cambridge University Press: Cambridge.

Friston, K. J., Harrison, L. and Penny, W. (2003) Dynamic causal modelling. NeuroImage 19, 1273– 1302.

Glasser, M. F., Sotiropoulos, S. N., Wilson, J. A., Coalson, T. S., Fischl, B., Andersson, J. L., Xu, J., Jbabdi, S., Webster, M., Polimeni, J. R., Van Essen, D. C., Jenkinson, M. and Consortium, W. U.-M. H. (2013) The minimal preprocessing pipelines for the Human Connectome Project. NeuroImage 80, 105–124.

Griffanti, L., Salimi-Khorshidi, G., Beckmann, C. F., Auerbach, E. J., Douaud, G., Sexton, C. E., Zsoldos, E., Ebmeier, K. P., Filippini, N., Mackay, C. E., Moeller, S., Xu, J., Yacoub, E., Baselli, G., Ugurbil, K., Miller, K. L. and Smith, S. M. (2014) ICA-based artefact removal and accelerated fMRI acquisition for improved resting state network imaging. NeuroImage 95, 232–247.

Hinojosa, C. A., George, G. C. and Ben-Zion, Z. (2024) Neuroimaging of posttraumatic stress disorder in adults and youth: progress over the last decade on three leading questions of the field. Molecular psychiatry 29, 3223–3244.

Hodgkin, A. L. and Huxley, A. F. (1952) A quantitative description of membrane current and its application to conduction and excitation in nerve. The Journal of physiology 117, 500–544.

Horn, A. and Blankenburg, F. (2016) Toward a standardized structural-functional group connectome in MNI space. NeuroImage 124, 310–322.

Horn, A., Neumann, W. J., Degen, K., Schneider, G. H. and Kuhn, A. A. (2017) Toward an electrophysiological “sweet spot” for deep brain stimulation in the subthalamic nucleus. Hum. Brain Mapp. 38, 3377–3390.

Jirsa, V., Wang, H., Triebkorn, P., Hashemi, M., Jha, J., Gonzalez-Martinez, J., Guye, M., Makhalova, J. and Bartolomei, F. (2023) Personalised virtual brain models in epilepsy. The Lancet Neurology 22, 443–454.

Kawamura, Y., Nakao, H. and Kuramoto, Y. (2007) Noise-induced turbulence in nonlocally coupled oscillators. Physical review. E, Statistical, nonlinear, and soft matter physics 75, 036209.

Kolmogorov, A. N. (1941a) Dissipation of Energy in Locally Isotropic Turbulence. Proceedings of the USSR Academy of Sciences (in Russian) 32, 16–18.

Kolmogorov, A. N. (1941b) The local structure of turbulence in incompressible viscous fluid for very large Reynolds numbers. Proceedings of the USSR Academy of Sciences (Atmos. Ocean. Phys.) 30, 299–303.

Kuramoto, Y. (1984) Chemical Oscillations,Waves, and Turbulence. Springer-Verlag: Berlin.

Lorenz, E. N. (1993) The essence of chaos. University of Washington Press: Seattle.

Martinez-Molina, N., Sanz-Perl, Y., Escrichs, A., Kringelbach, M. L. and Deco, G. (2024) Turbulent dynamics and whole-brain modeling: toward new clinical applications for traumatic brain injury. Front Neuroinform 18, 1382372.

Navarro Schroder, T., Haak, K. V., Zaragoza Jimenez, N. I., Beckmann, C. F. and Doeller, C. F. (2015) Functional topography of the human entorhinal cortex. eLife 4.

Oostenveld, R., Fries, P., Maris, E. and Schoffelen, J. M. (2011) FieldTrip: Open source software for advanced analysis of MEG, EEG, and invasive electrophysiological data. Comput. Intell. Neurosci. 2011, 156869.

Ott, E. and Antonsen, T. M. (2008) Low dimensional behavior of large systems of globally coupled oscillators. Chaos: An Interdisciplinary Journal of Nonlinear Science 18, 037113.

Perl, Y. S., Mininni, P., Tagliazucchi, E., Kringelbach, M. L. and Deco, G. (2023a) Scaling of whole-brain dynamics reproduced by high-order moments of turbulence indicators. Physical Review Research 5, 033183.

Perl, Y. S., Zamora-Lopez, G., Montbrió, E., Monge-Asensio, M., Vohryzek, J., Fittipaldi, S., Campo, C. G., Moguilner, S., Ibañez, A., Tagliazucchi, E., Yeo, B. T. T., Kringelbach, M. L. and Deco, G. (2023b) The impact of regional heterogeneity in whole-brain dynamics in the presence of oscillations. Network neuroscience 7, 632–660.

Piccinini, J., Deco, G., Kringelbach, M., Laufs, H., Sanz Perl, Y. and Tagliazucchi, E. (2022) Data-driven discovery of canonical large-scale brain dynamics. Cereb Cortex Commun 3, tgac045.

Ponce-Alvarez, A., Deco, G., Hagmann, P., Romani, G. L., Mantini, D. and Corbetta, M. (2015) Resting-state temporal synchronization networks emerge from connectivity topology and heterogeneity. PLoS Comput. Biol. 11, e1004100.

Protas, B. (2008) Vortex dynamics models in flow control problems. Nonlinearity 21, R203.

Saffman, P. G. (1992) Vortex dynamics. In: Theoretical Approaches to Turbulence. pp. 263–277. Springer.

Salimi-Khorshidi, G., Douaud, G., Beckmann, C. F., Glasser, M. F., Griffanti, L. and Smith, S. M. (2014) Automatic denoising of functional MRI data: combining independent component analysis and hierarchical fusion of classifiers. NeuroImage 90, 449–468.

Schaefer, A., Kong, R., Gordon, E. M., Laumann, T. O., Zuo, X. N., Holmes, A. J., Eickhoff, S. B. and Yeo, B. T. T. (2018) Local-Global Parcellation of the Human Cerebral Cortex from Intrinsic Functional Connectivity MRI. Cereb. Cortex 28, 3095–3114.

Setsompop, K., Kimmlingen, R., Eberlein, E., Witzel, T., Cohen-Adad, J., McNab, J. A., Keil, B., Tisdall, M. D., Hoecht, P., Dietz, P., Cauley, S. F., Tountcheva, V., Matschl, V., Lenz, V. H., Heberlein, K., Potthast, A., Thein, H., Van Horn, J., Toga, A., Schmitt, F., Lehne, D., Rosen, B. R., Wedeen, V. and Wald, L. L. (2013) Pushing the limits of in vivo diffusion MRI for the Human Connectome Project. NeuroImage 80, 220–233.

Sip, V., Hashemi, M., Dickscheid, T., Amunts, K., Petkoski, S. and Jirsa, V. Characterization of regional differences in resting-state fMRI with a data-driven network model of brain dynamics. Science advances 9, eabq7547.

Smith, S. M., Beckmann, C. F., Andersson, J., Auerbach, E. J., Bijsterbosch, J., Douaud, G., Duff, E., Feinberg, D. A., Griffanti, L., Harms, M. P., Kelly, M., Laumann, T., Miller, K. L., Moeller, S., Petersen, S., Power, J., Salimi-Khorshidi, G., Snyder, A. Z., Vu, A. T., Woolrich, M. W., Xu, J., Yacoub, E., Ugurbil, K., Van Essen, D. C., Glasser, M. F. and Consortium, W. U.-M. H. (2013) Resting-state fMRI in the Human Connectome Project. NeuroImage 80, 144–168.

Snyder, J., Zlotnik, A. and Lokhov, A. Y. (2020) Data-driven selection of coarse-grained models of coupled oscillators. Physical Review Research 2, 043402.

Xu, Y., Long, X., Feng, J. and Gong, P. (2023) Interacting spiral wave patterns underlie complex brain dynamics and are related to cognitive processing. Nat Hum Behav 7, 1196–1215.

